# Application of a human lectin array to rapid *in vitro* screening of sugars used as targeting tags for therapeutics

**DOI:** 10.1101/2025.02.12.637837

**Authors:** Stefi V. Benjamin, Maureen E. Taylor, Kurt Drickamer

## Abstract

An increasing number of clinical applications employ oligosaccharides as tags to direct therapeutic proteins and RNA molecules to specific target cells. Current applications are focused on endocytic receptors that result in cellular uptake, but additional applications of sugar-based targeting in signaling and protein degradation are emerging. These approaches all require development of ligands that bind selectively to specific sugar-binding receptors, known as lectins. In the work reported here, a human lectin array has been employed as a predictor of targeting specificity of different oligosaccharide ligands and as a rapid *in vitro* screen to identify candidate targeting ligands. The approach has been validated with existing targeting ligands, such as a GalNAc cluster ligand that targets siRNA molecules to hepatocytes through the asialoglycoprotein receptor. Additional small oligosaccharides that can selectively target other classes of cells have also been identified and the potential of larger glycans derived from glycoproteins has been investigated. In initial screens, ligands for targeting either vascular or sinusoidal endothelial cells and plasmacytoid dendritic cells have been identified. Lectin array screening has also been used to characterize the specificity of glycolipid-containing liposomes that are used as carriers for targeted delivery. The availability of a rapid *in vitro* screening approach to characterizing natural oligosaccharides and glycomimetic compounds has the potential to facilitate selection of appropriate targeting tags before undertaking more complex *in vivo* studies.

## Introduction

Use of oligosaccharides (glycans) to target proteins, lipids and RNA molecules to specific tissues and cells has become a viable clinical approach relatively recently. Glycan targeting is mediated by various glycan-binding receptors (lectins) on the surfaces of specific cell types. For example, in enzyme replacement therapy for lysosomal storage disorders, enzymes are tagged with sugars that direct them to the mannose receptor on macrophages or to the mannose 6-phosphate receptor found on a wide range of cells [Grabowski et al. 1995; Lee et al. 2003]. More recently, siRNA treatments to reduce expression of blood proteins produced in the liver have been directed to the asialoglycoprotein receptor on hepatocytes with a glycan tag [Nair et al. 2014; Brannagan et al. 2022]. The tag used in these cases is a cluster of GalNAc residues identified through a series of ligand-optimization studies [Prakash et al. 2016].

Targeting liposomes through glycan tags has been investigated *in vitro* using GM1, GM3 and synthetic sialoside ligands that interact with sialoadhesin (siglec-1/CD169) or other siglecs [Affandi et al. 2020; Shen et al. 2024] and there have been some early-stage clinical applications [Arena et al. 2022]. In addition to these applications that have reached the clinic, other potential targeting strategies using glycan have been proposed. For example, antibodies tagged with sugar epitopes, known as lysosome-targeting chimeras (LYTACs) can potentially clear specific proteins from cell surfaces and from circulation [Banik et al. 2020; Ahn et al. 2021; Wang et al. 2024]. In all of these cases, successful targeting depends on selective binding of the glycan tag to an appropriate receptor and significant *in vivo* screening is required in the design of appropriate glycan conjugates.

Mammalian lectin arrays have recently been developed for rapid screening of glycoconjugate binding to a panel of sugar-binding receptors [Jégouzo et al. 2020; Benjamin et al. 2024]. These arrays can be used to test binding of glycans on bacteria, viruses, fungi and parasites in order to investigate how these micro-organisms interact with cells in the innate immune system. In addition to demonstrating the roles of these receptor in pathogen recognition, availability of a comprehensive panel of human sugar-binding receptors could also facilitate rapid *in vitro* screening of glycans and glycomimetics to assess their potential for *in vivo* targeting.

The utility of the human lectin array for characterizing the interactions of existing targeting glycans has now been demonstrated and additional oligosaccharides that can potentially be used as receptor- and cell-specific delivery tags have been identified.

## Results

### Strategy

The current version of the human lectin array displays 39 different carbohydrate-recognition domains (CRDs) from 36 receptors, covering 7 different structural categories of lectins (Table 1). Several approaches were used to test the ability of oligosaccharide epitopes to target specific receptors (Figure 1). The primary strategy was to use biotinylated glycans complexed with fluorescently-labelled streptavidin for initial screening. Screening was repeated at a 25-to 100-fold range of concentrations. In most cases, the results were similar across concentrations, suggesting that binding is relatively high affinity, but in a few cases binding to some receptors decreases with reduced ligand concentration, consistent with weaker binding. Ligands that showed selective binding in the tetravalent format could then be further tested with streptavidin modified so that it forms only monomers. In some cases, bivalent complexes were formed with antibodies to biotin and highly multivalent ligands were either serum albumin with covalently attached sugars or liposomes. Comparison of these results provides some insights into whether selective binding results from affinity for a single oligosaccharide or requires multivalent presentation.

**Table 1.**
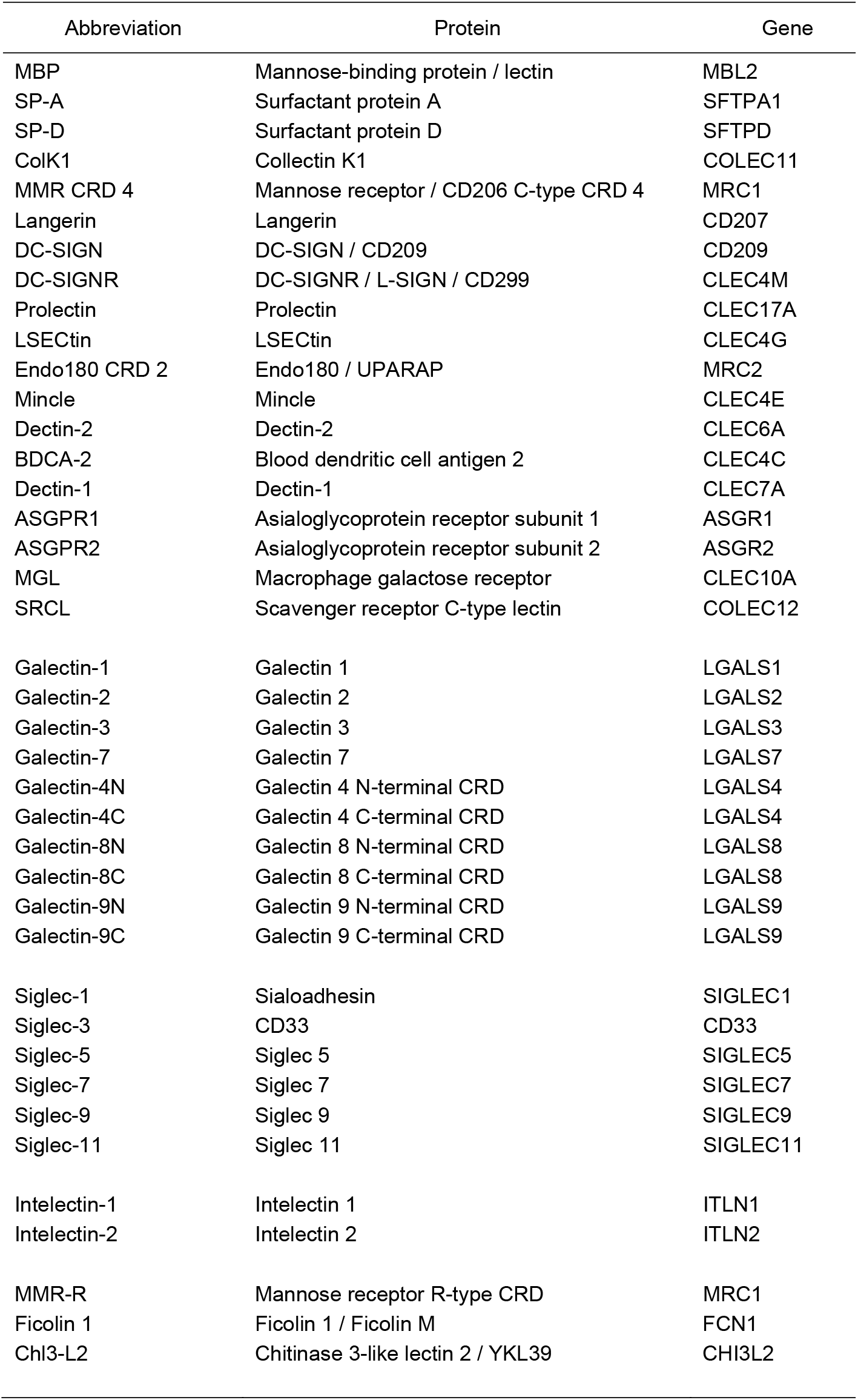
CRDs displayed on human lectin array.

**Figure 1.**
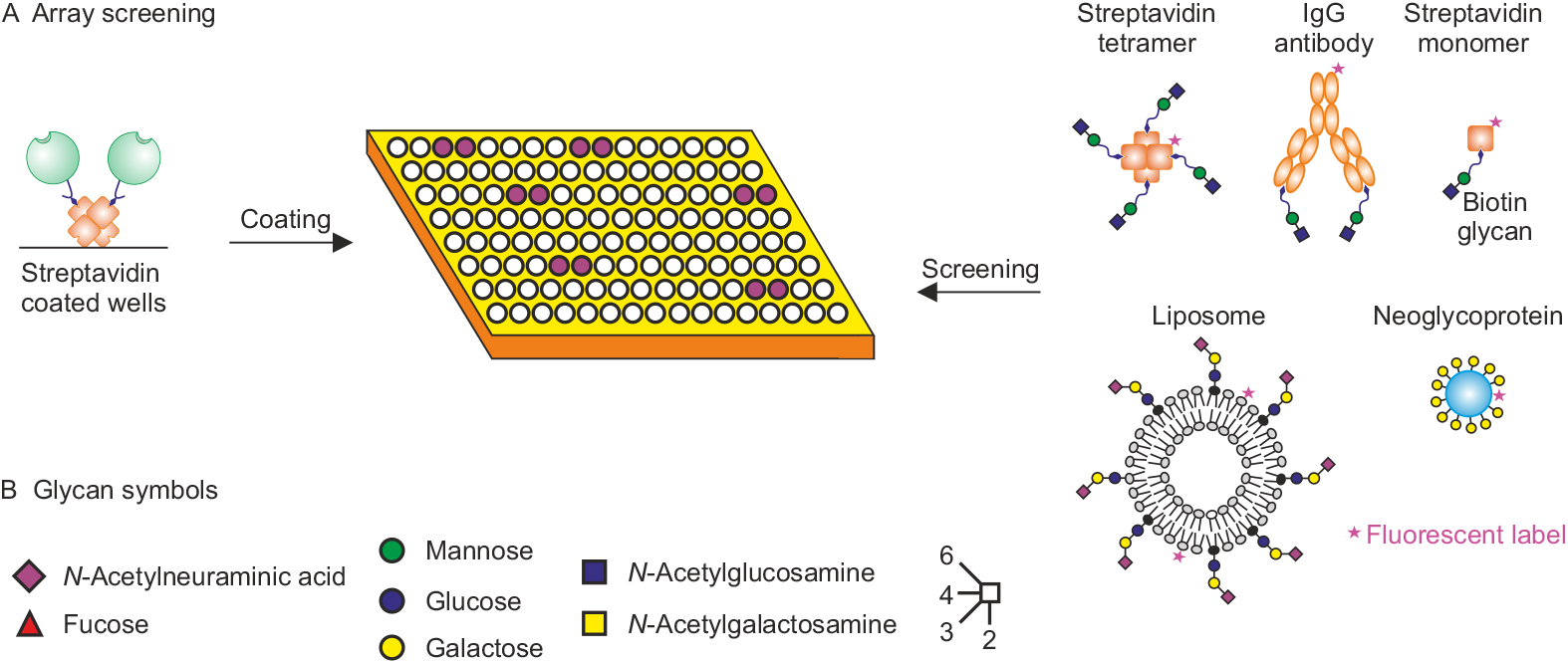
Strategy for screening of lectin array with receptor-specific epitopes. (A) Methods for presentation of glycans as multivalent, tetravalent, bivalent and monovalent ligands are summarized. (B) Glycan symbols used here and in subsequent figures. Unless otherwise indicated, linkages from galactose and GlcNAc residues are in β configuration and linkages from NeuAc, fucose, mannose, and GalNAc are in α configuration.

### Monosaccharides versus oligosaccharides

Screening the array with simple monosaccharide ligands, such as biotinylated galactose, complexed with streptavidin failed to show consistent binding to any receptors. In contrast, screening with the highly multivalent neoglycoprotein ligands Man_31_-BSA and Gal_33_-BSA in each case showed binding to multiple CRDs (Figure 2), confirming that the monosaccharides bind, but must be highly multivalent to achieve sufficient avidity to withstand washing of the array. While the binding shows clear selectivity of mannose and galactose for different classes of CRDs, it also demonstrates that effective targeting requires more complex oligosaccharide ligands.

**Figure 2.**
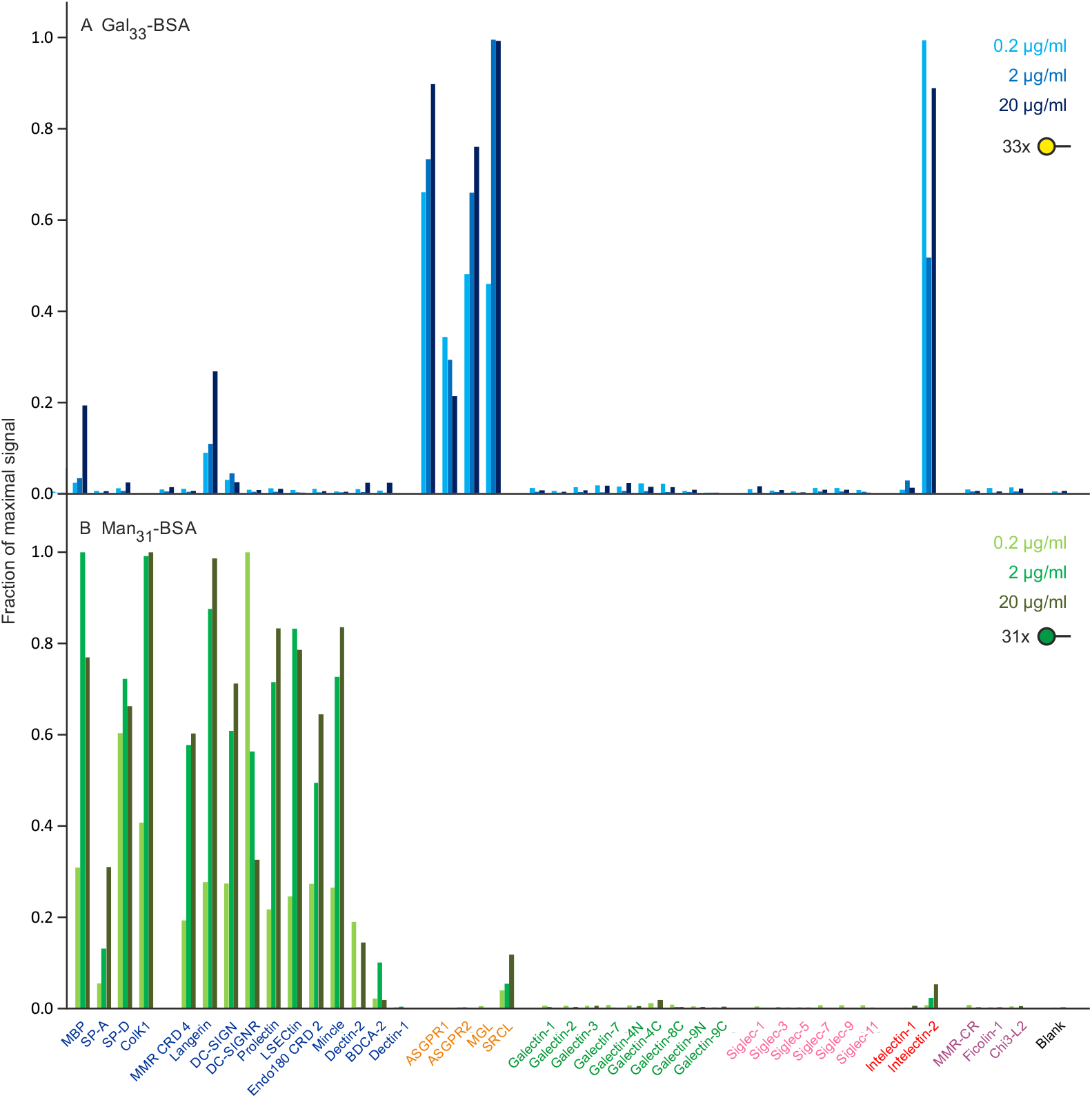
Screening of lectin array with FITC-labelled neoglycoproteins. (A) Gal_33_-BSA. Ranges for errors ranged from 11-13%. (B) Man_31_-BSA. Errors ranged from 15-15% (Table S1).

A widely used targeting glycomimetic developed for directing siRNA molecules to hepatocytes through the asialoglycoprotein receptor contains a cluster of three GalNAc residues [Prakash et al. 2016; Brannagan et al. 2022]. Streptavidin tetramers complexed with a biotinylated form of this synthetic cluster ligand showed very selective binding, with the strongest signals observed for the major subunit of the asialoglycoprotein receptor over a range of concentrations (Figure 3A). However, the results also indicate that this ligand binds to the macrophage galactose receptor. A similar pattern of binding was also observed for a monomeric version of this ligand (Figure 3B), reflecting tight binding of the ligand through a high-affinity binding site. Interestingly, a Galα1-3Galβ1-4GlcNAc trisaccharide complexed with streptavidin shows high specificity for the asialoglycoprotein receptor without binding the macrophage receptor, although it does interact weakly with some of the galectins (Figure 3C). However, this oligosaccharide does not show selective binding as a bivalent antibody complex (Table S3).

**Figure 3.**
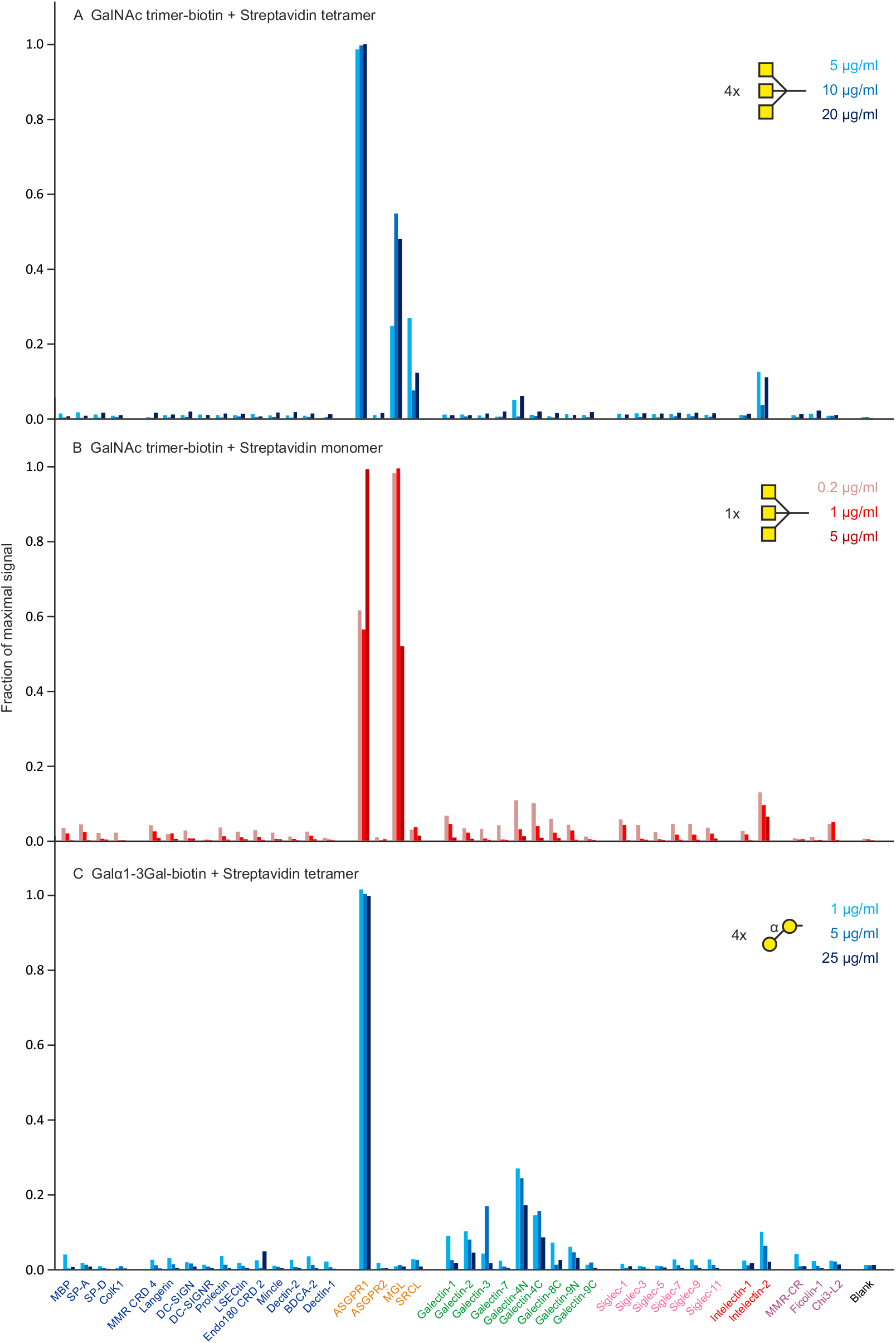
Screening of lectin array with GalNAc_3_ cluster ligand. (A) Cluster ligand complexed with streptavidin tetramer. Errors ranged from 11-15%. (B) Cluster ligand complexed with streptavidin monomer. Errors ranged from 2-16% (Tables S2 and S3).

The array results with the GalNAc cluster ligand correlate with observations in mice showing hepatic accumulation of siRNA conjugated to this ligand, which has been optimized for targeting the asialoglycoprotein receptor with a single glycan attached to one end of an RNA molecule [Prakash et al. 2016; Brannagan et al. 2022]. These experiments demonstrate the utility of the lectin array for demonstrating selective binding in a simple *in vitro* experiment. The observed binding to the macrophage galactose receptor also shows that binding to additional receptors may not always be detected in animal clearance studies, probably because of the relative abundance of the hepatic receptor.

### Identification of additional glycan epitopes with narrow receptor specificity

Screening with naturally occurring di- and trisaccharides revealed several additional candidates for selective receptor targeting. The targeting potential of these ligands depends both on a narrow receptor-binding profile for the glycan and on restricted expression of the receptor on specific cell types. Glycans that meet these criteria include the disaccharide GlcNAcβ1-2Man, which interacts almost exclusively with LSECtin, a receptor found only on sinusoidal endothelial cells [Powlesland et al. 2008; Liu et al. 2004], and the Lewis^x^ trisaccharide, which binds particularly well to the scavenger receptor C-type lectin (SRCL) that is found more generally on endothelial cells [Coombs et al 2005; Ohtani 2001; Graham et al. 2011].

Binding of GlcNAcβ1-2-Man to LSECtin is highly specific even as the valency is reduced from tetrameric streptavidin complexes (Figure 4A) to the bivalent antibody complexes (Figure 4B). Binding to surfactant protein SP-A was observed for all antibody complexes tested and reflects the interaction of the CRD from SP-A to the antibody Fc region in a carbohydrate-independent manner [Lin and Wright 2006]. Binding also remains largely specific with a monomeric complex (Figure 4C), although some binding to the B cell-specific receptor prolectin is observed.

**Figure 4.**
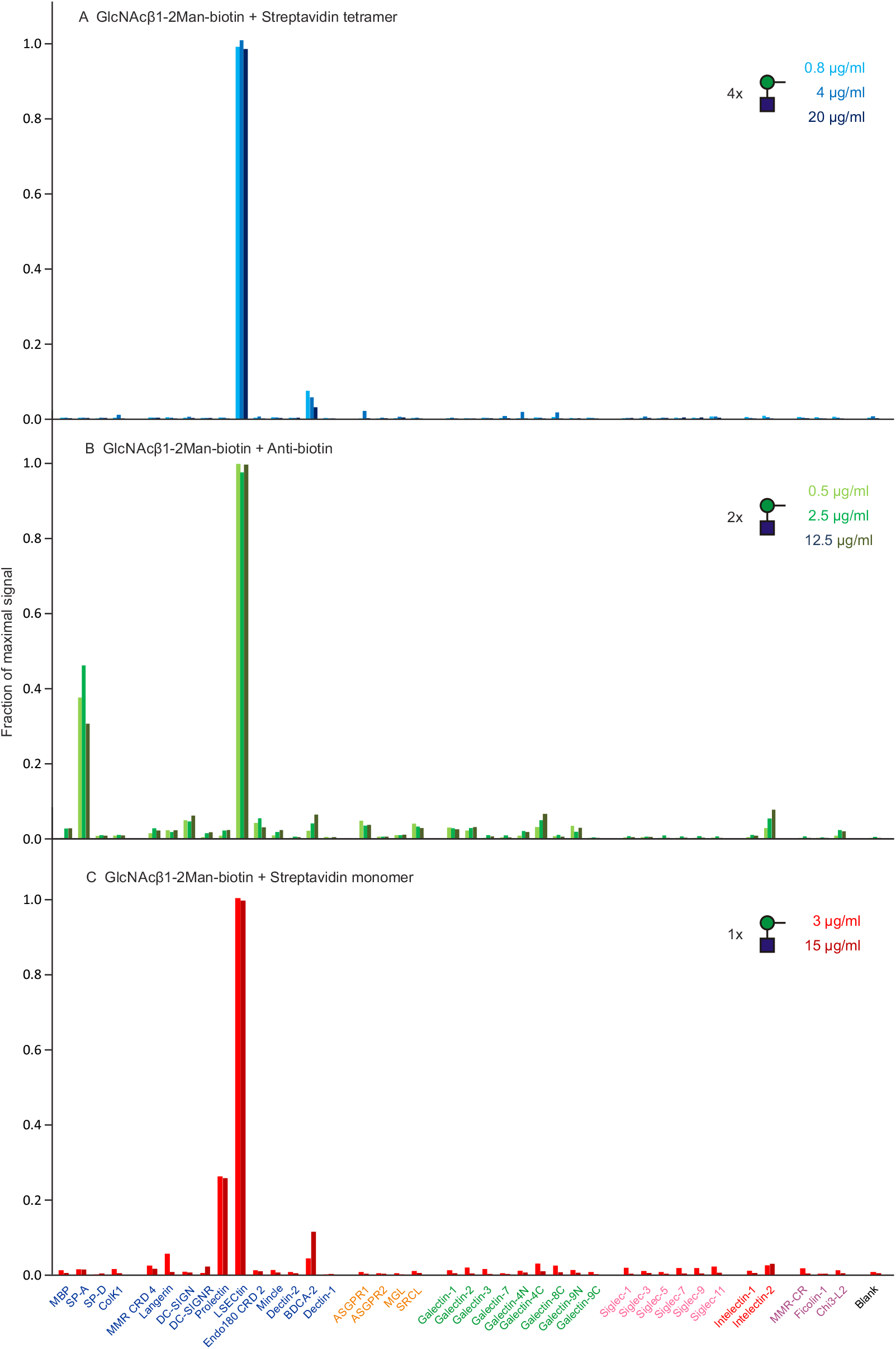
Screening of lectin array with GlcNAcβ1-2-Man disaccharide. (A) Complex with streptavidin tetramer. Errors ranged from 21-27%. (B) Complex with anti-biotin antibody. Errors ranged from 2-5%. (C) Complex with streptavidin monomer. Errors ranged from 12-18% (Table S4).

Binding of the Le^x^ trisaccharide Galβ1-3(Fucα1-4)GlcNAc shows high specificity for SRCL both as a tetrameric complex (Figure 5A) and as a bivalent antibody complex (Figure 5B). However, very little binding of a complex with streptavidin monomers was observed and it was not selective for SRCL (Table S5). Thus, the results for these two simple oligosaccharides suggest that selective binding to endothelial clearance receptors could be achieved with dimeric presentation of sugar epitopes.

**Figure 5.**
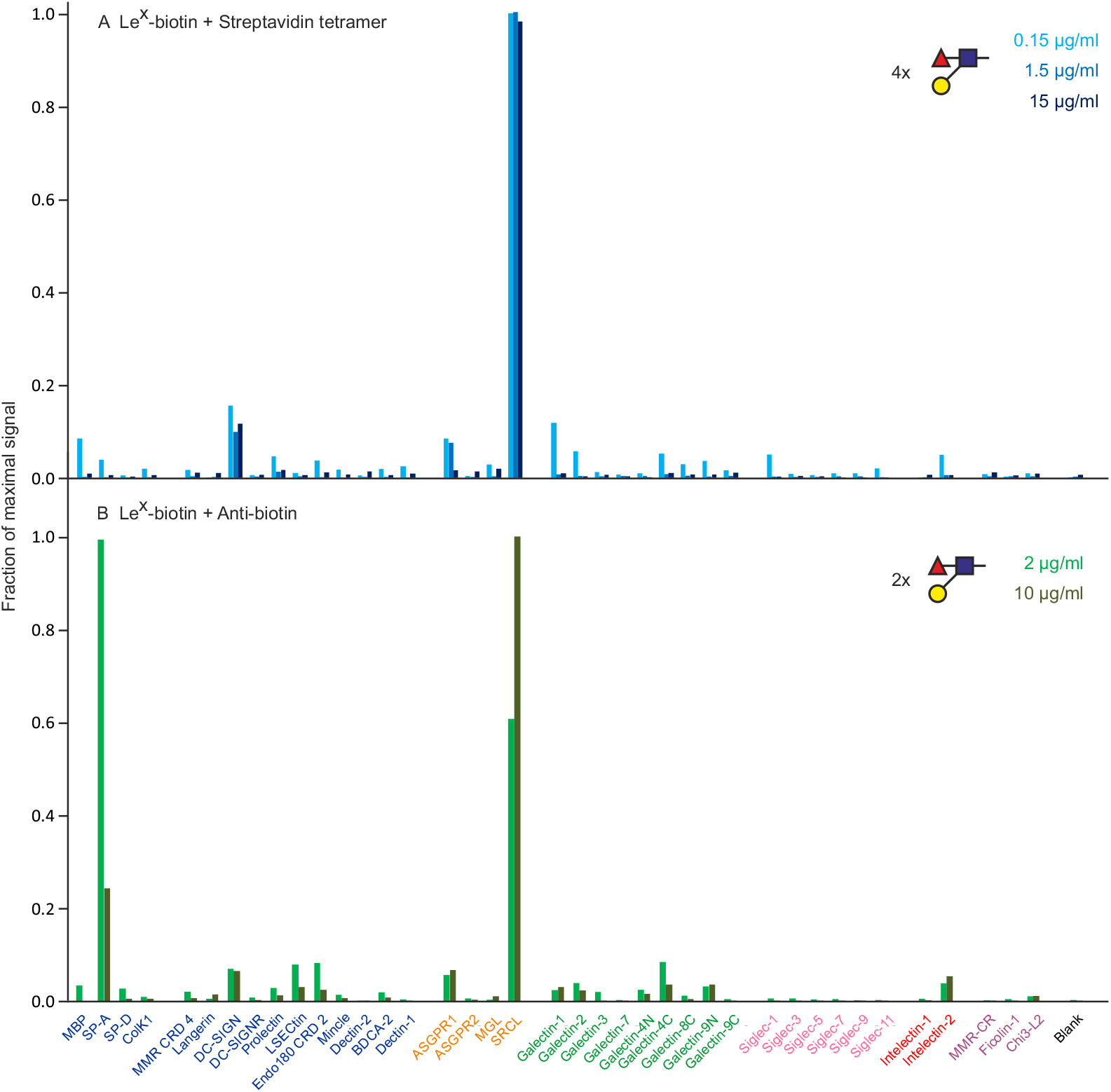
Screening of lectin array with Galβ1-3(Fucα1-4)GlcNAc (Le^X^) trisaccharide. (A) Complex with streptavidin tetramer. Errors ranged from 4-24%. (B) Complex with anti-biotin antibodies. Errors ranged from 6-9% (Table S5).

A tetrameric complex of the T-antigen disaccharide Galβ1-3GalNAc-α-largely targets galectin-4, through both the N- and C-terminal CRDs and preferential binding to galectin-4 is also evident in a monomeric complex (Figure 6). In this case, a multivalent neoglycoprotein was also available and showed binding to galectin-4, but also binding to the N-terminal CRD of galectin-9 and to the asialoglycoprotein receptor. Galectin-4 is expressed almost exclusively in the gut, so in this environment the interaction would likely be largely specific. Given the involvement of galectin-4 in tumor metastasis in the digestive system, the result suggests that inhibitory oligosaccharides based on the T-antigen could selectively block key interactions [Huflejt and Leffler 2004].

**Figure 6.**
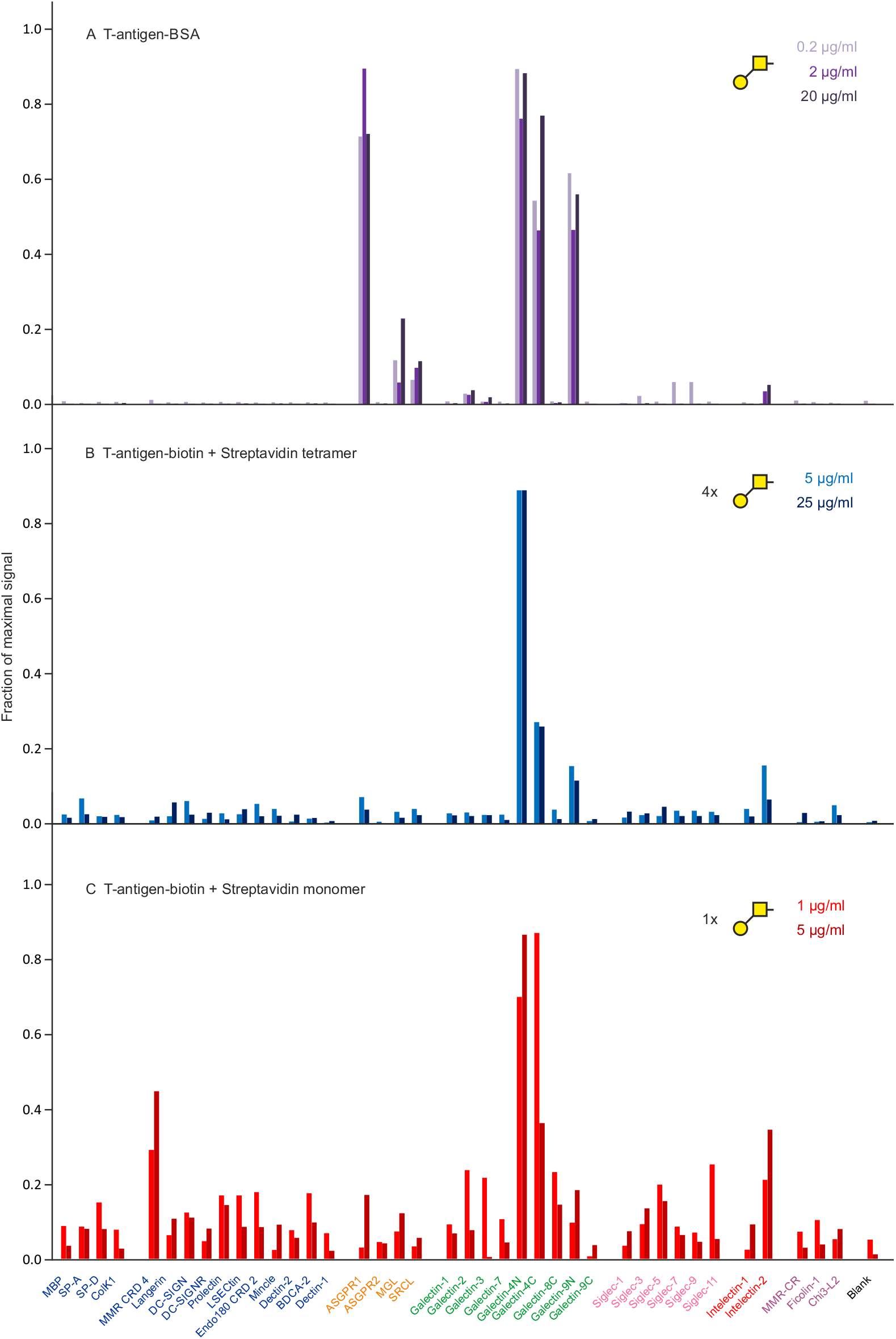
Screening of lectin array with Galβ1-3GalNAc-α-(T-antigen) disaccharide. (A) FITC-labelled T-antigen-BSA neoglycoprotein. Ranges for errors ranged from 6-9%. (B) Complex of oligosaccharide with streptavidin tetramer. Errors ranged from 7-28%. (C) Complex of oligosaccharide with streptavidin monomer. Errors ranged from 17-30% (Table S6).

### Screening with natural glycans

In addition to small oligosaccharides, some larger N-linked glycans also show significant selectivity for specific receptors. The two N-glycosylation sites in human transferrin are predominantly occupied by biantennary, complex glycans, with a very small amount of more branched structures [Yamashita et al. 1989]. Screening the array with desialylated transferrin showed binding to the dendritic cell antigen-2 (BDCA-2), a receptor that is expressed almost exclusively on plasmacytoid dendritic cells (Figure 7A) [Dzionek et al. 2001]. This binding is consistent with the binding characteristics of BDCA-2, which interacts predominantly with exposed Galβ1-4GlcNAc termini of these glycans [Jégouzo et al. 2015]. Binding to other CRDs on the array, including the asialoglycoprotein receptor and some of the galectins, is very concentration-dependent, suggesting that binding to these CRDs reflects lower affinity interactions.

**Figure 7.**
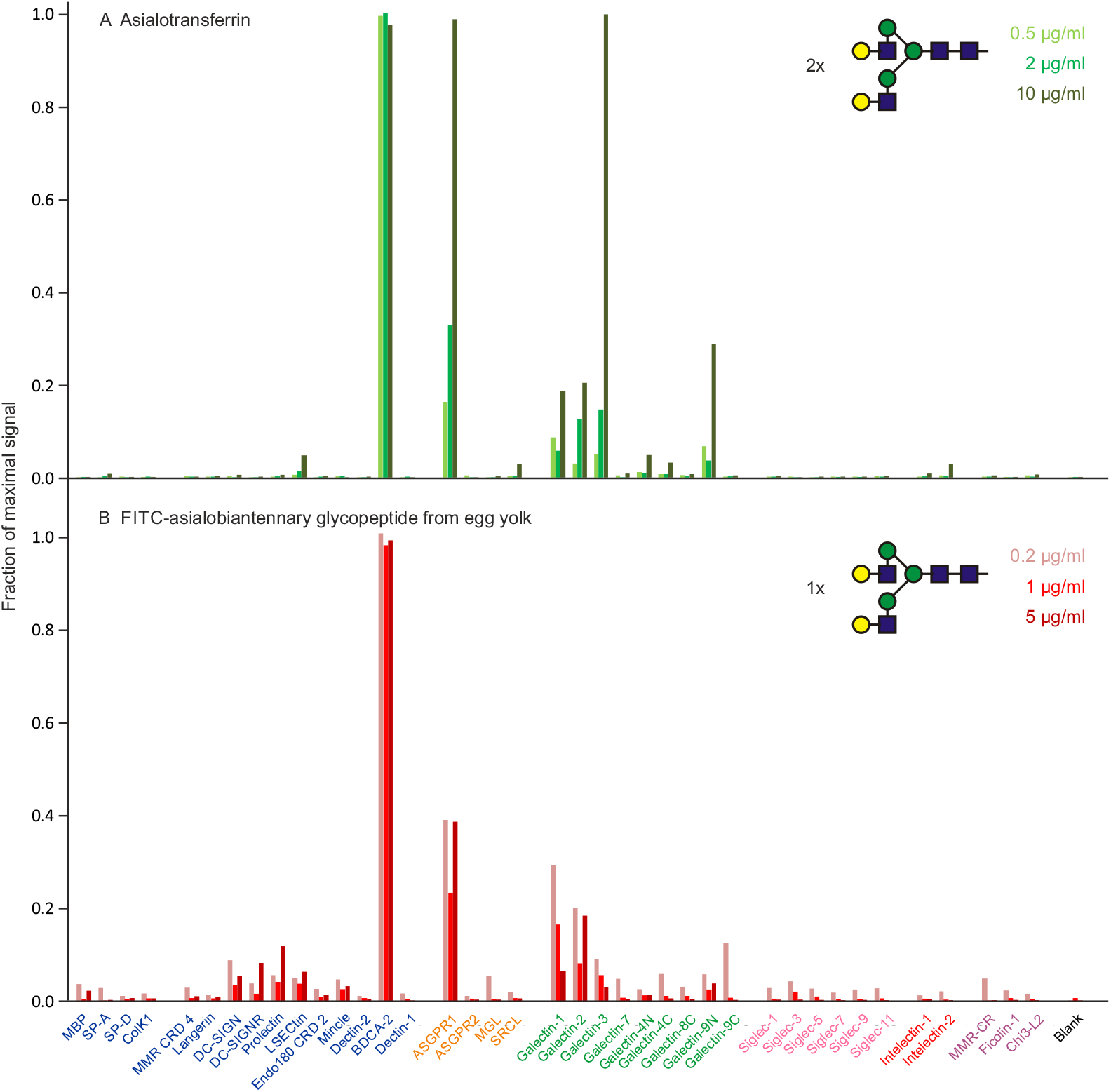
Screening of lectin array with Galβ1-4GlcNAc (LacNAc) disaccharide-containing conjugates of complex N-linked glycans. (A) FITC-labeled asialotransferrin. Ranges for errors ranged from 4-25%. (B) FITC-labelled egg oligosaccharide. Ranges for errors ranged from 10-14% (Table S7).

In order to confirm that binding to BDCA-2 results from the presence of biantennary glycans on asialotransferrin, an equivalent glycan generated from chicken egg yolk glycopeptide was directly labelled with FITC and used to screen the array (Figure 7B). The binding profile largely mirrors what was observed for asialo-transferrin. The low level of binding to several CRDs such as DC-SIGN, DC-SIGNR, prolectin, LSECtin and others, may result from the presence of a small fraction of glycans lacking a terminal galactose residue on one branch [Liu et al. 2017], thus leaving an exposed terminal GlcNAc residue that can bind to these CRDs. Although the N-linked glycans in transferrin are closely spaced and might both interact with immobilized CRDs, binding of the monomeric glycan suggests that such multivalent interaction is not necessary to achieve specificity. The very cell-type specific expression of BDCA-2 and the role of blood dendritic cells in regulating production of interferon α make this a potentially useful targeting interaction [Dzionek et al. 2001].

In contrast to very selective binding of the complex N-linked glycans, oligomannose-type N-linked glycans interact with a wide range of CRDs (Figure 8). Both Man_5_- and Man_9_-containing glycans bind most effectively to the dendritic cell receptor DC-SIGN, but both also bind to multiple other receptors. Thus, while these oligosaccharides have shown some potential for targeting to dendritic cells through DC-SIGN [Valverde et al. 2020], it may prove difficult to make this receptor a specific target when all of the other potential targets are considered.

**Figure 8.**
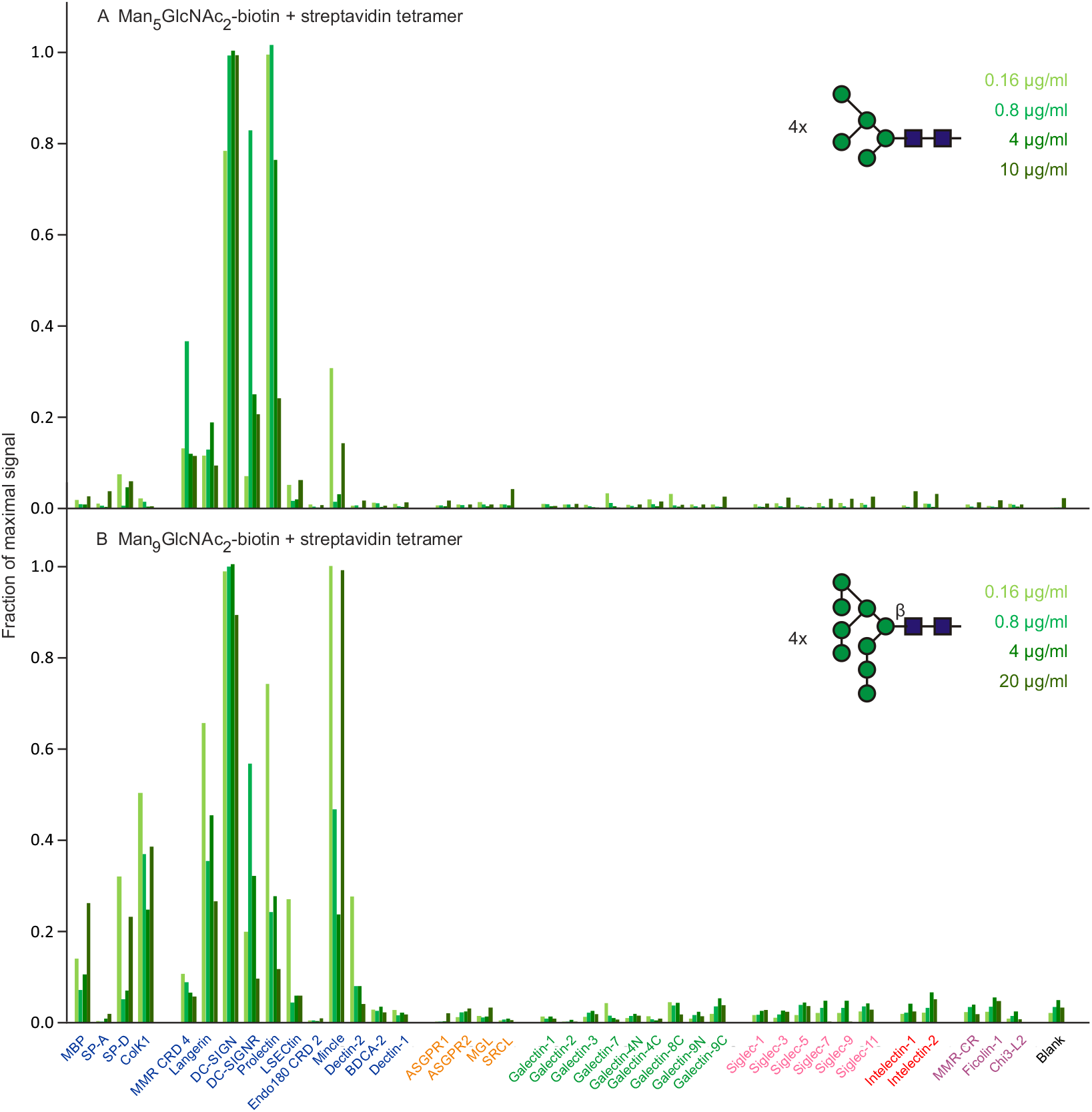
Screening of lectin array with oligomannose-containing glycan ligands. (A) Man_5_GlcNAc_2_. Errors ranged from 4-15%. (B) Man_9_GlcNAc_2_. Errors ranged from 11-17% (Table S8).

The Man_5_ glycan is somewhat more selective than Man_9_, particularly because it shows a much reduced signal for CRDs from the soluble serum collectins. The results suggest that the terminal Manα1-2Man caps on the Man_9_ glycan may be responsible for the additional interactions, although screening with short Manα1-2Man polymers on their own did not demonstrate selective targeting (Table S8).

### Screening with liposomes

Liposomes containing either GM1 or GM3 gangliosides have been used to target the sialic-binding receptor sialoadhesin on macrophages (Siglec-1; CD169) [Affandi et al. 2020; Shen et al. 2024]. Screening of the lectin array with fluorescently labelled liposomes containing these gangliosides confirmed that both show strong binding to sialoadhesin (Figure 9). Amongst the siglecs tested, GM1-containing liposomes are selective for sialoadhesin, while those containing GM3 interact with siglec 3 (CD33) to a similar extent and more weakly with other siglecs. Thus, in terms of targeting siglecs, GM1-containing liposomes are potentially more specific.

**Figure 9.**
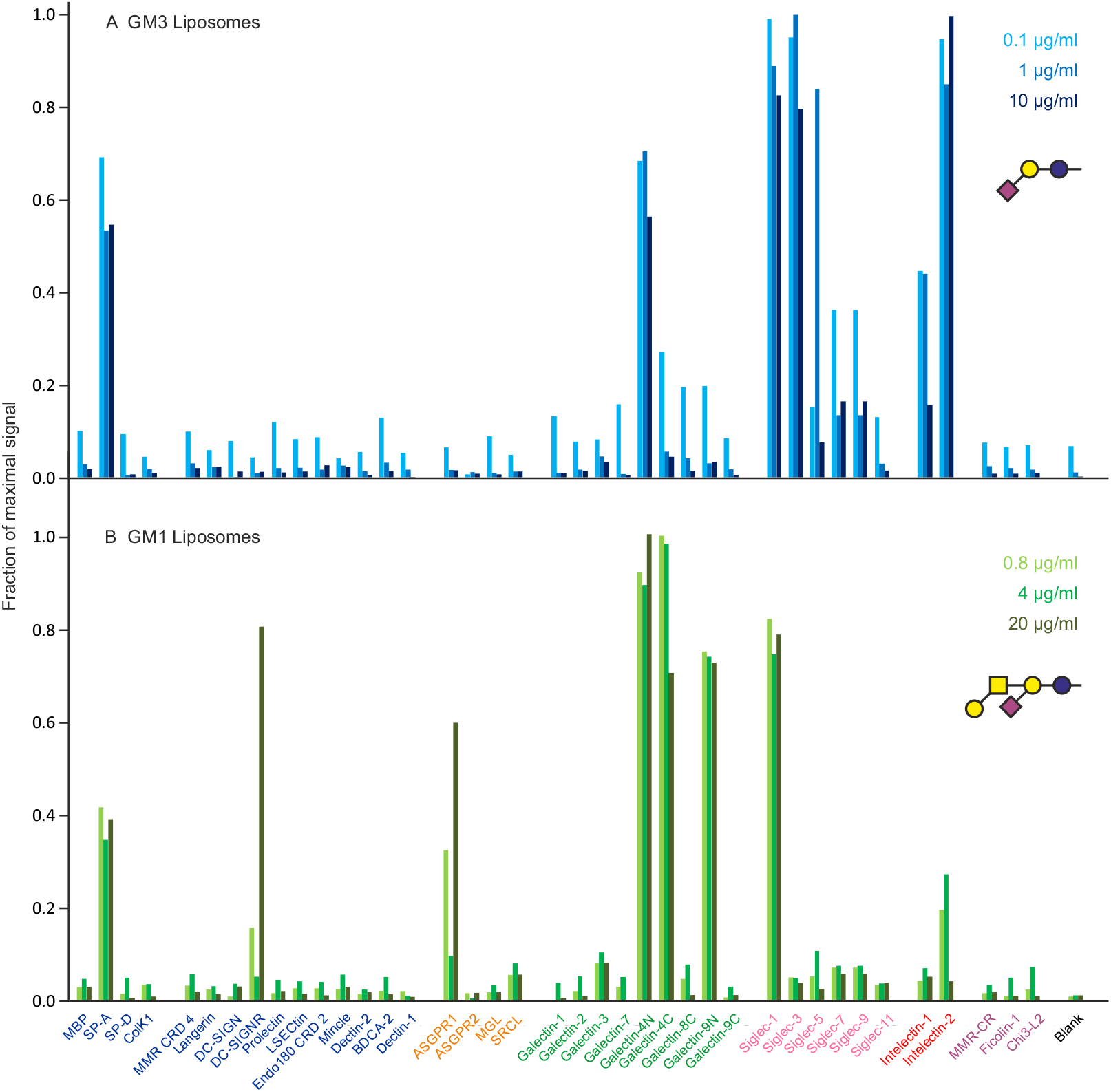
Screening of lectin array with ganglioside-containing liposomes. (A) GM1. Errors ranged from 7-10%. (B) GM3 (Table S9).

Both types of liposomes interact with the N-terminal CRD from galectin-4, but the GM1 liposomes also bind to the C-terminal CRD of galectin 4 and the N-terminal CRD of galectin 9. This binding pattern is reminiscent of the binding of T-antigen to these same galectin CRDs (Figure 6), reflecting the presence of the Galβ1-3GalNAc structure in the headgroup of GM1. GM1 also binds to intelectin-2, probably through the galactose residue (unpublished observations). Binding to the CRD from pulmonary surfactant protein SP-A is observed for all liposome preparations tested, including those lacking glycans, and thus likely reflects the well characterized capacity of this CRD to bind lipids [McCormack et al. 1997]. Given the distinct sites of expression of the siglecs, largely in the immune system, and galectins 4 and 9 as well as intelectin-2 in the gut and other epithelia, targeting of injected liposomes would presumably be primarily dictated by the different siglec-binding properties of the gangliosides [Huflejt and Leffler 2004; Nonnecke et al. 2022; Gonzalez-Gil and Schnaar 2021].

## Discussion

Table 2 summarizes the binding interactions reported here, along with information about the localization of some of the human glycan-binding receptors that can be selectively targeted by oligosaccharide ligands. The results indicate that this screening can provide useful guidance for development of sugar-based targeting tags. A key outcome is identification of some simple oligosaccharides that show strong selectivity for specific receptors that have restricted tissue distributions.

**Table 2.**
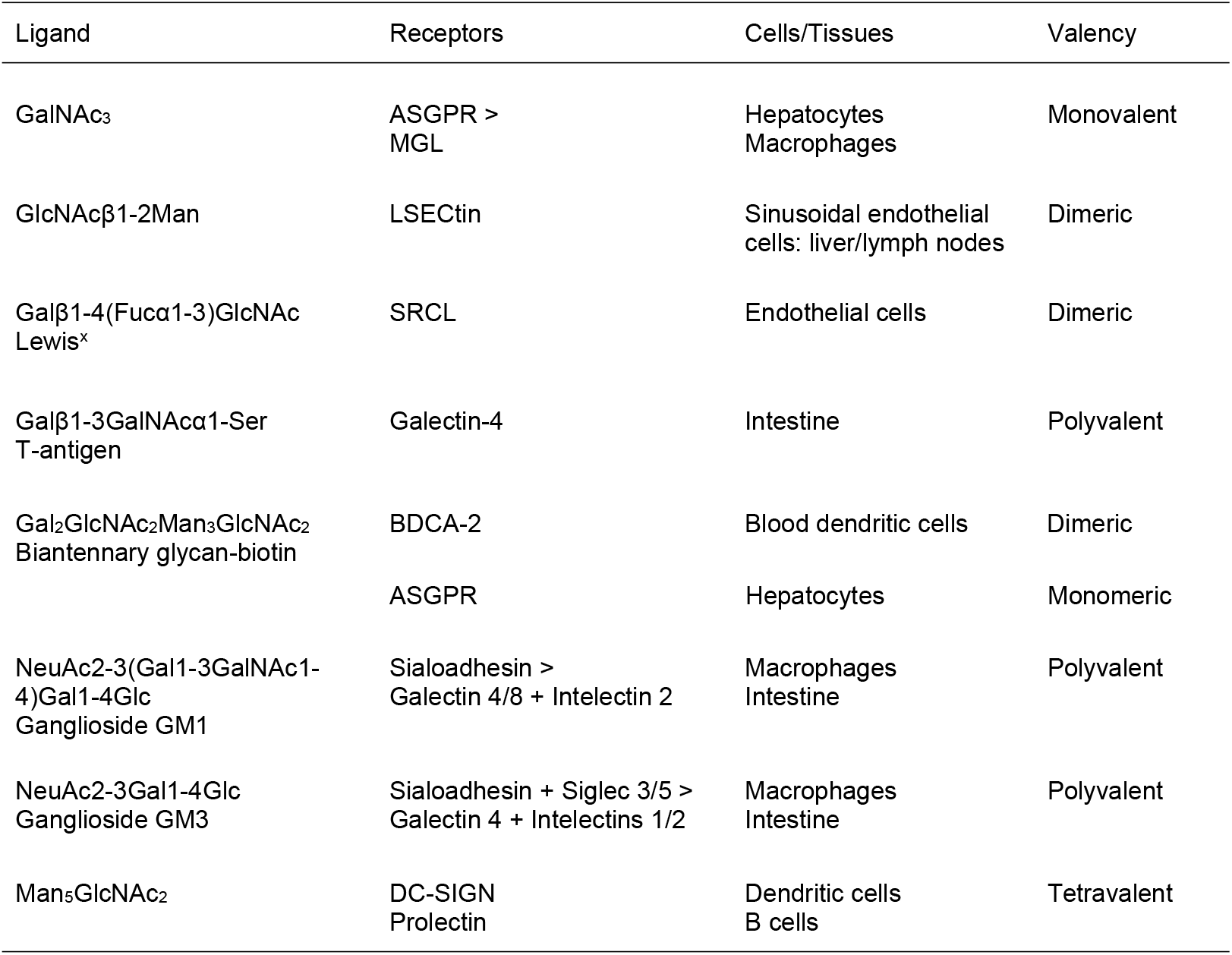
Summary of selective ligand-receptor interactions Valency indicates the lowest valency of ligand that shows selective binding.

| Ligand | Receptors | Cells/Tissues | Valency |
| --- | --- | --- | --- |
| GalNAc<sub>3</sub> | ASGPR ><br>MGL | Hepatocytes<br>Macrophages | Monovalent |
| GlcNAcβ1-2Man | LSEctin | Sinusoidal endothelial<br>cells: liver/lymph nodes | Dimeric |
| Galβ1-4(Fucα1-3)GlcNAc<br>Lewis<sup>x</sup> | SRCL | Endothelial cells | Dimeric |
| Galβ1-3GalNAcα1-Ser<br>T-antigen | Galectin-4 | Intestine | Polyvalent |
| Gal<sub>2</sub>GlcNAc<sub>2</sub>Man<sub>3</sub>GlcNAc<sub>2</sub><br>Biantennary glycan-biotin | BDCA-2 | Blood dendritic cells | Dimeric |
|  | ASGPR | Hepatocytes | Monomeric |
| NeuAc2-3(Gal1-3GalNAc1-4)Gal1-4Glc<br>Ganglioside GM1 | Sialoadhesin ><br>Galectin 4/8 + Intelectin 2 | Macrophages<br>Intestine | Polyvalent |
| NeuAc2-3Gal1-4Glc<br>Ganglioside GM3 | Sialoadhesin + Siglec 3/5 ><br>Galectin 4 + Intelectins 1/2 | Macrophages<br>Intestine | Polyvalent |
| Man<sub>5</sub>GlcNAc<sub>2</sub> | DC-SIGN<br>Prolectin | Dendritic cells<br>B cells | Tetravalent |

The positive results suggest that it may be possible to identify additional oligosaccharides with unique targeting potential. One potential candidate is GalNAc-4-SO_4_, which interacts with the R-type CRD in the macrophage mannose receptor [Fiete et al. 1998]. Looking beyond mammalian glycans, microbial glycans may provide inspiration for other useful sugar epitopes. For example, although the mammalian mannose-terminated glycans show a disappointing lack of specificity, substructures found in yeast mannan, such as Manα1-2Man oligomers, might show greater selectivity for receptors such as dectin-2 [Feinberg et al. 2017]. Similarly, GlcNAc-containing polymers such as poly-N-acetylglucosamine (GlcNAcβ1-4GlcNAc) found in biofilms shows more restricted binding [Benjamin et al. 2024], suggesting other types of oligosaccharides that might be tested.

Successful development of the synthetic GalNAc cluster ligand demonstrates that improvements in affinity and cost can be achieved using glycomimetics [Prakash et al. 2016]. Such synthetic analogs can be optimized by monitoring affinity for a proposed target receptor, but monitoring specificity using the array yields complementary information, by providing a rapid *in vitro* method for screening out ligands that are less specific for the target. Obtaining this information in an *in vitro* assay could reduce the need for more complex studies in animals during the initial screening steps.

Screening the array can also flag up issues that may reduce targeting specificity *in vivo*. The results confirm specific targeting by the GalNAc cluster ligand that is already employed in several approved clinical treatments that direct siRNA molecules to hepatocytes. Although the data suggest that some of this ligand will be directed to macrophages, the high capacity of the hepatic clearance system would mean that most of the siRNA would be directed to hepatocytes [Nair et al. 2014]. The Galα1-3Galβ1-4GlcNAc trisaccharide shows higher specificity for the asialoglycoprotein receptor without binding the macrophage receptor, but does not work as a monomeric ligand.

The results for desialylated egg glycopeptide suggest that even limited heterogeneity of natural glycans can results in some mis-targeting, although the resulting loss of specificity could be prevented with appropriate quality control. The liposome screening experiments also demonstrate that in addition to binding to sialoadhesin, the intended target ligand on macrophages, natural gangliosides will likely also target both other siglecs and lectins in other families. Development of glycomimetics specific for individual siglec addresses the first point [Nycholat et al. 2019], but screening against the larger complement of CRDs on the array would potentially provide further evidence of specificity.

The results presented here are largely focused on oligosaccharide tags that target receptors for uptake into cells. However, receptor-specific oligosaccharides identified using the array can be employed in other applications, such as triggering signaling through receptors that activate intracellular kinases and phosphatases. Combining the specificity results from the array with other information about receptor geometry can provide a basis for designing novel stimulatory ligands. For example, the demonstration that bi-antennary, galactose-terminated glycans bind preferentially to the BDCA-2 receptor on plasmacytoid dendritic cells, combined with structural information about the arrangement of BDCA-2 dimers, suggests ways to create ligands that could modulate interferon α production by forming BDCA-2 clusters [Liu et al. 2024].

## Materials and Methods

### Glycoproteins

For fluorescein labelling, 1 mg of neoglycoproteins Gal_33_-BSA or Man_31_-BSA from E-Y laboratories was reacted with 12.5 μg FITC in 250 μl of 100 mM bicine, pH 9.0 for 2 h at room temperature. Excess reagent was removed by repeated washing with Tris-buffered saline in a VivaSpin-2 centrifugal concentrator with a 10-kDa cut-off membrane (VIVAproducts). Human transferrin (1 mg) from Sigma was digested with 1250 units of *Clostridium perfringens* neuraminidase from New England Biolabs in 50 μl of 50 mM sodium acetate buffer, pH 5.5, 5 mM CaCl_2_, for 20 h at 37 °C. The sample was diluted into 350 mM bicine, pH 9.0 and labelled as described above.

### Egg glycopeptide

Egg glycopeptide, prepared and desialylated as described previously [Jégouzo et al. 2015], was labelled with FITC and purified on a 1 × 30 cm BioGel P2 column run in Tris-buffered saline.

### Liposomes

Liposomes were prepared by combining 3 μmole of distearoyl phosphatidylcholine, 1.25 μmole of M1 or GM3 gangliosides (Avanti Polar Lipids), 1.75 μmole of cholesterol, and 0.25 μg of aminofluorescein coupled to distearoyl phosphatidyl ethanolamine-polyethylene glycol 2000-N-hydroxysuccinimide (Cayman Chemicals). The mixture was dried, resuspended in 2 ml of TBS, sonicated for 1 min and extruded 5 times through 0.2 μm aluminum filters (Anitop).

### Complexes with biotinylated oligosaccharides

Biotinylated oligosaccharides GlcNAcβ1-2Man and Le^x^ were from Dextra Laboratories and T-antigen, Man_5_GlcNAc_2_ and GalNAc_3_ cluster ligand were from Sussex Research Laboratory. 1,2-α-1,2-α-D-mannotriose-1-O-ethylamine from Biosynth was reacted with biotinamido-hexanoic acid *N*-hydroxysulfosuccinimide ester from Sigma. The product was purified by chromatography on BioGel P2 and Dowex-1 columns and characterized by mass spectrometry. AlexaFluor488-labelled streptavidin was from Life Technologies and AlexaFluor488-labelled anti-biotin was from Jackson Immunoresearch. Monomeric streptavidin from Sigma was labelled with FITC as described above. Biotinylated oligosaccharides were combined with streptavidin or antibody in a 10-fold molar excess compared to biotin-binding sites. Complexes were formed by incubation for 3 h at room temperature in Tris-buffered saline, pH 7.4, and were purified on a 15-ml column of Sephadex G-25 eluted with Tris-buffered saline.

### Array screening

All procedures were conducted in binding buffer containing 0.15 M NaCl, 25 mM Tris-Cl, pH 7.8, 2.5 mM CaCl_2_ in streptavidin-coated 96-well black plates (Life Technologies). Coating of wells with biotinylated CRDs was performed as previously described [Benjamin et al. 2024]. Ligands diluted in binding buffer containing 0.1% bovine serum albumin were added in 60 µl aliquots. For complexes of biotinylated ligands, binding was performed in the presence of 20 mM free biotin. After incubation for 2-4 h at 4 °C, wells were washed 3 times with binding buffer and scanned directly on a Victor3 (PerkinElmer) or a ClarioStar (BMG Labtech) multiwell plate reader.

In all cases, averages of duplicate wells are plotted, with error bars representing the range of values. In each experiment, results are normalized to the maximum signal. For each ligand used to screen the array, average percentage errors given in the legends were determined as the difference between the values for duplicate wells as a percentage of the average of the values. The overall average errors for each ligand were based on signals that were greater than 10% of the maximum signal.

## Supporting information

Supporting Information cover page

## Data availability

All data are contained in the manuscript and supporting information.

## Supporting information

This article contains supporting information.

## Author contributions

M. E. T., K. D. conceptualization; S. V. B., M. E. T., K. D. investigation; S. V. B., M. E. T., K. D. data curation; M. E. T., K. D. writing - original draft; S. V. B., M. E. T., K. D. writing - review & editing; S. V. B., K. D. visualization.

## Funding and additional information

This work was supported by UKRI Biotechnology and Biological Sciences Research Council grant BB/V014137/1 to K. D. and M. E. T.

## Abbreviations

The abbreviations used are:

CRD: carbohydrate-recognition domain
FITC: fluorescein isothiocyanate
BSA: bovine serum albumin
SRCL: scavenger receptor C-type lectin
BDCA-2: blood dendritic cell antigen 2
DC-SIGN: dendritic cell-specific intercellular adhesion molecule 1-grabbing nonintegrin.

